# Comparative Analysis of Soil Microbial Communities in High-Tunnel and Field Agricultural Systems

**DOI:** 10.64898/2026.02.06.704486

**Authors:** Mark H. Timper, Daniel Schlatter, Natalie Hoidal, Devanshi Khokhani

## Abstract

High tunnels and open-field systems differ markedly in soil physicochemical properties, yet their effects on belowground microbiomes remain poorly understood. We characterized bacterial and fungal communities in paired high-tunnel and adjacent field soils from 100 small-scale vegetable farms across Minnesota, integrating amplicon sequencing of 16S rRNA and ITS2 regions with soil nutrient data, arbuscular mycorrhizal fungi (AMF) spore counts, and microbial co-occurrence networks. High-tunnel soils had higher pH, organic matter, and multiple macronutrients (notably P, K, and N forms) and lower bulk density than fields, reflecting intensive organic amendments and reduced leaching. Despite these differences, bacterial and fungal alpha diversity did not differ between environments, whereas beta diversity analyses revealed strong shifts in community composition. High tunnels were enriched in salt- and stress-tolerant bacterial phyla (Firmicutes, Deinococcota, Patescibacteria, Halanaerobiaeota, Halobacterota) and saprotrophic fungal groups (Mortierellomycota, Ascomycota, Basidiomycota, Mucoromycota), while several oligotrophic or symbiotic taxa, including Acidobacteriota and Glomeromycota, declined. Glomeromycota relative abundance was negatively correlated with high soil phosphorus, whereas AMF spore densities did not decline, suggesting suppression of active mycorrhizal symbioses rather than propagule loss under high-nutrient conditions. Co-occurrence network analyses showed that bacterial and fungal networks in high tunnels were less dense, more modular, and exhibited higher ratios of positive to negative associations than field networks, consistent with stress-induced shifts toward more facilitative interactions. Collectively, our results indicate that high-tunnel production homogenizes soil microbiomes and selects for stress- and high-nutrient-adapted taxa, with potential consequences for nutrient cycling, AMF function, and long-term agroecosystem outcomes.

## Introduction

Soil microbiomes, complex communities of bacteria, fungi, archaea, microscopic protists, and viruses, are a critical part of terrestrial ecosystems. These microorganisms interact with one another and with the environment, facilitating organic matter decomposition and nutrient cycling while indirectly maintaining plant health (Schulz et al. 2013; Fierer 2017). Their activity contributes to soil fertility, structure, water retention, and disease resistance, making them a crucial component of sustainable agriculture (Hartmann and Six 2022; Suman et al. 2022). As agriculture adapts to rapidly and continually fluctuating weather patterns and a growing world population, understanding how management practices influence soil microbial communities is essential to the development of productive and resilient agroecosystems.

Agricultural management practices, including conventional and organic farming, tillage, and monoculture/intercropping, are known to impact soil microbial abundance and composition significantly (García-Orenes et al. 2013; Hartmann et al. 2015; Mondaca et al. 2024). These shifts arise from management-induced changes in soil structure, moisture, aeration, and nutrient availability. Soil microorganisms are key drivers of soil health; therefore, choosing management practices that support microbial diversity for activities such as nutrient cycling and overall plant health, or those that have less impact on microbial diversity, is crucial for the long-term sustainability of agroecosystems (Gomiero et al. 2011). High tunnels, also known as hoop houses, are semi-controlled environments that are increasingly used to extend growing seasons and protect crops from environmental stressors (Lamont 2009; Kandel et al. 2020). Unlike greenhouses, high tunnels typically lack automated climate control but are cheaper to build and maintain, especially for small-scale growers. In Minnesota, where weather extremes can significantly impact growing seasons, high tunnels have become increasingly popular. Natural Resources Conservation Service (NRCS) High Tunnel Initiative funded around 700 high tunnels in MN from 2010-2020 (Pierre et al. 2024). Despite their benefits, high tunnels may alter soil conditions, affecting microbial communities by altering temperature, moisture, and nutrient availability (Bruce et al. 2019). Recent research by (Hoidal et al. 2026) collected and analyzed soil samples from 100 Minnesota farms, finding that high-tunnel soils had significantly higher levels of organic matter, pH, nitrate, ammonium, phosphorus, and potassium than field soils.

These elevated nutrient levels are thought to result from excessive nutrient inputs of compost and composted manure. Further, unique soil microclimates, including areas within high-tunnels that receive no irrigation, may also contribute to differences in nutrient cycling. Additionally, these high tunnels receive no water during the winter months, unlike fields, which receive snow and rain. Given the sensitivity of soil microbes to physicochemical properties, these differences could influence microbial diversity and community structure (Zhang et al. 2023). Soil nutrient availability also impacts the composition and functional traits of soil microbiomes, which are crucial for nutrient cycling and maintaining sustainable soil health (Zhang et al. 2024; Paes da Costa et al. 2024). Previous studies have investigated the individual effects of different management practices and these soil properties on soil microbial communities, but there is limited research on how these affect microorganisms in high tunnels compared to field environments. It is essential to address this knowledge gap as high tunnels become increasingly prevalent, enabling farmers to adopt sustainable farming practices and potentially enhance crop productivity. This study attempts to address these knowledge gaps by building on results from Hoidal et al (Hoidal et al. 2026) and addressing key questions regarding the effects of management practices and soil properties on soil microorganisms.

Using DNA sequencing of bacterial and fungal marker genes, we assessed microbial diversity and community composition of soils collected from field and high tunnels from 100 locations across Minnesota (Fig. 1). Given the uneven distribution of soil moisture, with dripline areas having more water than areas farther from the dripline, we collected samples from both areas within the high tunnel. The primary research questions we asked were: (1) Are there differences in soil microbial alpha and beta diversity between high tunnel and field environments? (2) Do specific soil nutrients such as phosphorus, potassium, nitrate, and ammonium, or pH and organic matter, have a significant impact on soil microbial beta diversity across and within high-tunnel and field environments? (3) Do microbial networks have distinct structures in high-tunnel and field environments?

**Figure 1.**
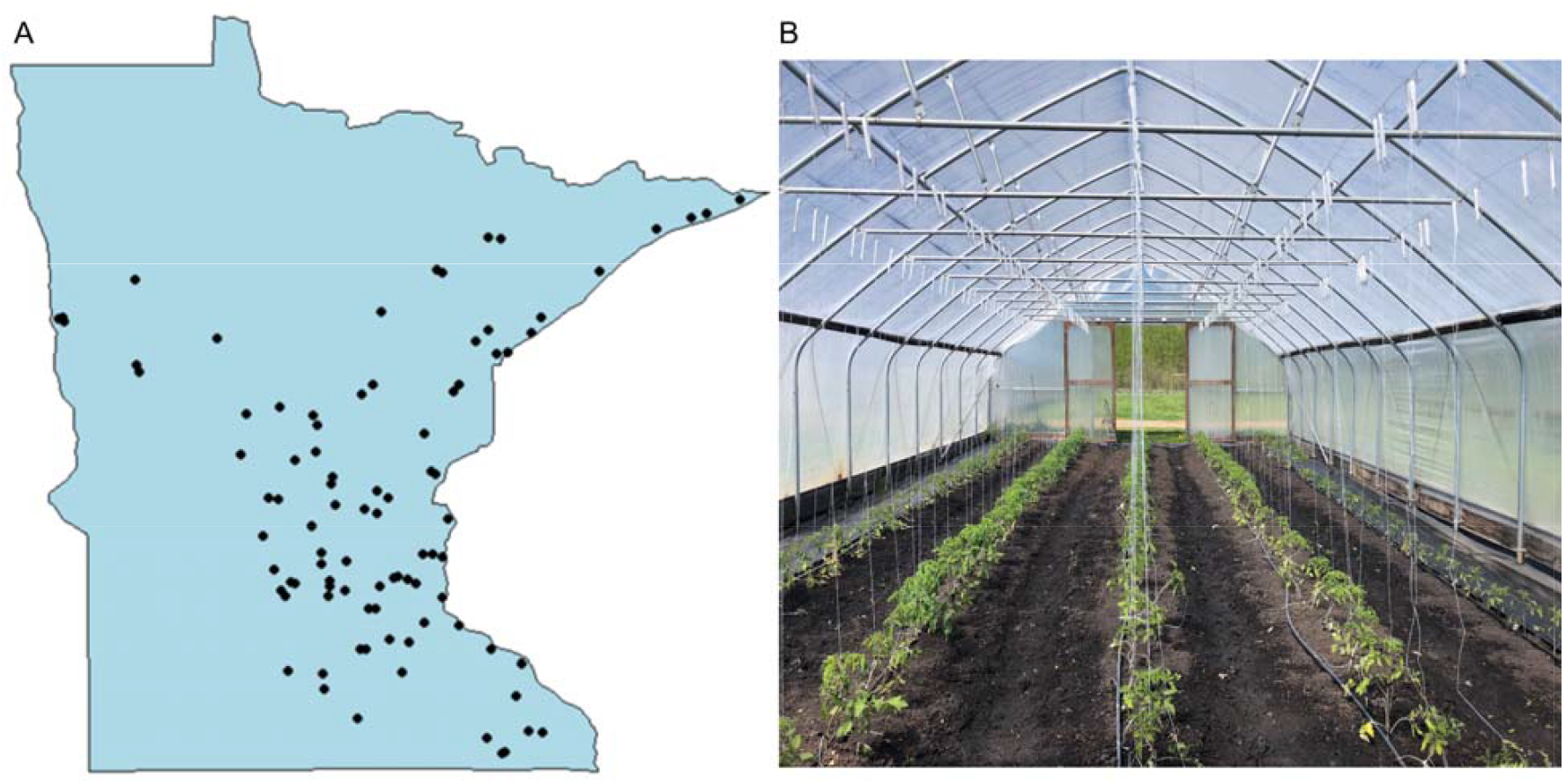
Soil collection sites. A) Map of Minnesota indicating sites from where the high-tunnel and field soil samples were collected. B) Image of one of the high tunnels to show sites away and close to irrigation lines.

## Materials and Methods

### Soil Collection and Testing

In spring 2023, soil samples from high tunnels and fields were collected from 100 small-scale vegetable farms in Minnesota by local extension educators (Hoidal et al. 2025). Each participating farm completed a survey detailing management and site history for their field and high tunnel plots. Sampling locations were pre-selected to represent the top 20 and bottom 20 values for phosphorus, pH, organic matter, nitrate, ammonium, and potassium. Soil samples were sent to the University of Minnesota Research Analytical Laboratory for analysis to measure their physicochemical properties. Each soil sample was briefly blended (pulsed) for up to 15 seconds in a Hamilton Beach Single Serve Blender. A no.35 sieve was stacked on a no.500 sieve, and the blended soil was passed through the no.35 sieve.

The blender and sieves were sterilized with 85% ethanol in between soil samples. The fine soil was poured into its corresponding bag, and 0.250 ± 0.005 grams of the soil was weighed and added to a clean 1.5 ml microcentrifuge tube. The 1.5 ml tubes containing soil were stored at - 20°C until needed.

### DNA extractions

DNA extractions were performed using the DNeasy PowerSoil Pro Kit from QIAGEN on 209 soil samples. As a negative control, 250 mL nuclease-free water was used instead of 250 mg of soil, to ensure that there was no contamination in the reagents. As a positive control, DNA from a mock microbial community (ZymoBIOMICS) was used. DNA quality was measured using a NanoDrop, checking the 260/280 ratio to ensure the purity of the extractions.

### DNA Sequencing

DNA extractions were sent to the University of Minnesota Genomic Center (UMGC) for AVITI 2×300 sequencing. The V3-V4 region of 16S rRNA gene (bacterial) and ITS2 region of the rRNA ITS region (fungal) were amplified and sequenced per UMGC protocols (Gohl et al. 2016a,b). Include UMGC protocol here. Briefly, primary PCRs using the respective primer sets (V3F: 5’CCTACGGGAGGCAGCAG-3’ and V4R: 5’-GGACTACHVGGGTWTCTAAT-3’ for the V3-V4 region and 5.8SF 5’-TCGATGAAGAACGCAGCG-3’ and ITS4R 5’-TCCTCCGCTTATTGATATGC-3’ for the ITS2 region) including Nextera adapters were performed using an initial denaturation step at 95C for 5 min, followed by 25 cycles of 98 °C for 20s, 55 °C for 15s, and 72 °C for 1 min, with a final extension at 72 °C for 5 min. These products were diluted 1:100, and 5 μL was used in a second PCR using forward (5′-AATGATACG GCGACCACCGAGATCTACAC[i5]TCGTCGGCAGCGTC-3′) and reverse (5′-CAAGCAGAAGACGGCA TACGAGAT[i7] GTCTCGTGGGCTCGG-3′) indexing primers.

This PCR consisted of an initial denaturation step at 95 °C for 5 min, followed by 10 cycles of 98 °C for 20s, 55 °C for 15s, and 72 °C for 1 min, with a final extension step at 72 °C for 5 min.

These products were pooled prior to sequencing. Raw fastq files are available under the NCBI SRA under the accession PRJNA1405668.

### R

All data processing and statistical analyses were performed in R (version 4.2.0 or 4.4.1). The packages ape, Biostrings, cowplot, dada2, devtools, ggplot2, ggplotify, igraph, Matrix, phyloseq, readr, ShortRead, SpiecEasi, tidyverse, viridis, and vegan were used to create and analyze results.

### Pre-Processing Data

Raw sequencing data from UMGC were processed using the DADA2 pipeline to remove low-quality reads, correct errors, and generate amplicon sequence variant (ASV) tables (Callahan et al. 2016). Briefly, primers were trimmed, low-quality ends truncated, and filtered reads were error-corrected to delineate ASVs. Cutadapt (Martin 2011) was used to trim fungal ITS2 reads prior to denoising to eliminate read-through. Chimeric sequences were filtered and taxonomy was assigned to remaining ASVs using the SILVA database (version 138.1) for bacteria and the UNITE (general release version 04.04.2024) for fungi. Once cleaned, ASV tables were rarefied using the phyloseq package in R. For the V3-V4 data, samples were excluded if they had less than 7,500 sequences, while for the ITS2 data, samples were excluded if they had less than 30,000 sequences. Rarefied data were used for all subsequent analyses unless otherwise specified.

### Data analysis

#### Alpha Diversity

The Shannon index (H’), Inverse Simpson index (1/D), and ASV richness, were used to assess the alpha diversity of the soil microbiome samples. These metrics were calculated using the vegan package in R. The Kruskal-Wallis test, a nonparametric test, was then performed to assess whether there were statistically significant differences in alpha diversity between field and tunnel environments across all metrics. Beta Diversity: Non-Metric Multidimensional Scaling (NMDS) on Bray-Curtis dissimilarity matrices was used to visualize beta diversity using the vegan package and plotted using ggplot2. To evaluate the relationship between soil properties and beta diversity, the envfit() function was used to fit vectors of soil properties to NMDS ordinations. Permutational multivariate analysis of variance (PERMANOVA) tests were performed using the adonis2() function to evaluate significant differences in microbial community composition among field and high-tunnel environments.

Differential abundance testing: Differentially abundant taxa among environments were assessed at the phylum and genus levels using ALDEx2 (Fernandes et al. 2014). Briefly, unrarefied ASV tables were agglomerated at the taxonomic level of interest and the ALDEx2 algorithm was used to contrast field vs. tunnel environments using the Kruskal-Wallis test of the ALDEx2 algorithm. Only taxa with Benjamini-Hochberg-corrected p-values <0.05 were considered significant. All samples from within tunnels were considered for these contrasts since comparisons of tunnel samples from close and away from driplines did not show any significant differences.

#### Co-occurrence Networks

Microbial co-occurrence networks were constructed for each environment. ASV tables were filtered to include only taxa present in >10% of samples and a mean relative abundance >0.05%. Spearman rank correlations were used to assess co-occurrence patterns using a threshold of rho > 0.6 and a Benjamini-Hochberg-corrected p-value of < 0.05 to define significant co-occurrence.

Global network statistics were calculated for each network, including the number of nodes, number of edges, graph density, mean degree, clustering coefficient (transitivity), modularity, number of positive edges, number of negative edges, and the positive: negative edge ratio using the igraph package in R. Networks were compared using Jaccard similarity.

### Arbuscular mycorrhizal fungal spore extraction

Arbuscular mycorrhizal fungal spores were extracted and enumerated using the wet sieving method (Gerdemann and Nicolson 1963), decanting, and followed by sucrose density gradient centrifugation. Briefly, 50 g of a soil sample was placed in a Waring Blender and blended at high speed for approximately 5 seconds to break up root fragments and release spores attached to roots or soil aggregates. The blended material was immediately poured through two sieves: the top sieve (500 µm openings), which captures roots and large debris, and the bottom sieve (38 µm openings), which captures most spores. The material from the bottom sieve was collected in a 50-mL beaker with a rubber policeman, then transferred into 50-mL tubes containing a 60% sucrose solution and water. These tubes were centrifuged (960 x g) for 5 minutes in a swinging-bucket rotor in a tabletop centrifuge. At the end of the run, the sucrose layer containing the spores was collected by pouring it onto a 38 µm sieve. These spores were washed with water before being collected onto petri plates using a rubber policeman. The bottoms of the plates were labeled with blue horizontal and red vertical lines to count the spores per square under the stereo microscope, and the number of spores per 50 g of soil sample was reported.

## Results

### Microbial diversity

Based on soil physicochemical properties reported in recent work (Hoidal et al. 2026), soil nutrients, including phosphorus, nitrate, potassium, calcium, magnesium, and sodium were all significantly higher in high tunnels than in open fields. Additionally, organic matter and cation exchange capacity were significantly higher in high tunnels than in open fields, and bulk density was significantly lower. Arbuscular mycorrhizal spore counts did not differ significantly between environments (Hoidal et al. 2026). Considering these observations, we examined the soil bacterial and fungal communities of samples with the highest and lowest values for the above-mentioned soil physicochemical properties (Table 1).

**Table 1:** Soil properties among field and high-tunnel environments. Different letters indicate significant differences between field and high-tunnel environments (Hoidal et al., 2026).

| Variable | Field | Tunnel |
| --- | --- | --- |
| Phos | 82.35 ± 73.86a | 169.78 ± 137.96b |
| OM | 4.6 ± 2.55a | 6.99 ± 3.59b |
| pH | 6.82 ± 0.72a | 7.16 ± 0.61b |
| NO<sub>3</sub> | 11.59 ± 11.38a | 73.29 ± 78.3b |
| NH<sub>4</sub> | 7.05 ± 5.65a | 9.4 ± 10.96a |
| K | 231.05 ± 206.52a | 360.88 ± 366.44b |
| CEC | 20.3 ± 13.58a | 29.66 ± 12.35b |
| Bulk Density | 1.15 ± 0.31a | 1.02 ± 0.27b |
| Ca | 3133.44 ± 2040.22a | 4394.75 ± 1799.38b |
| Mg | 453.94 ± 382.08a | 721.19 ± 367.41b |
| Na | 30.63 ± 28.93a | 106.52 ± 104.14b |
| Al | 0.64 ± 1.89a | 0.65 ± 1.27a |
| AMF Spore Count | 538.81 ± 403.69a | 640.93 ± 436.46a |

Values represent mean ± standard deviation. Metrics include amplicon sequence variants (ASV) richness, Shannon diversity (H’), and Simpson’s diversity (1/D). Significance was assessed using Kruskal-Wallis tests, and groups sharing the same letter did not differ significantly.

Alpha diversity metrics did not differ significantly among environments for either bacteria or fungi (Table 2). Bacterial ASV richness, Shannon diversity, and Simpson diversity were similar across field, tunnel-away, and tunnel-close soils, with no significant differences. Fungal alpha diversity showed similar patterns, with no significant differences among environments. These results indicate that high tunnel systems did not impact the overall microbial richness or within-sample diversity metrics for either bacteria or fungi.

**Table 2:** Alpha diversity of bacteria and fungi within fields and away from and close to drip irrigation in the high tunnels.

|  | Alpha Diversity | Field | Tunnel_Away | Tunnel_Close |
| --- | --- | --- | --- | --- |
| **Bacteria** | ASV Richness | 2792.97 ± 86.16a | 2771.49 ± 114.44a | 2884.31 ± 88.97a |
|  | Shannon H' | 7.29 ± 0.06a | 7.24 ± 0.07a | 7.31 ± 0.07a |
|  | Simpsons 1/D | 818.1 ± 29.38a | 760.9 ± 36.87a | 820.84 ± 30.79a |
| **Fungi** | ASV Richness | 534.01 ± 18.1a | 554.94 ± 19.88a | 573.57 ± 17.86a |
|  | Shannon H' | 4.59 ± 0.04a | 4.65 ± 0.05a | 4.66 ± 0.05a |
|  | Simpsons 1/D | 36.49 ± 1.92a | 39.25 ± 2.14a | 37.52 ± 2.08a |

Bacterial and fungal community composition differed significantly between environments. Bacterial communities (Fig. 2A) and fungal communities (Fig. 2C) each showed distinct separation between field and high-tunnel soils, though there was a large amount of variation across samples from each environment. However, community composition did not differ between dripline positions, with samples collected close to or away from the dripline tending to cluster together. PERMANOVA results showed no significant differences in bacterial or fungal community composition between samples collected away from and close to the dripline (p=1.0 for both groups). In contrast, community composition differed significantly between tunnel-close and field samples for both bacteria and fungi (p=0.0001). Vector fitting identified soil properties that were significantly correlated with community structure. For both bacteria (Fig. 2B) and fungi (Fig. 2D), vectors for CEC, pH, Mg, Ca, NO_3_ ^-^, Na, K, and P tended to be associated with high tunnel samples, reflecting the higher levels of many of these soil nutrients measured in tunnel samples compared to field samples. In contrast, greater soil bulk density tended to be associated with microbial communities from field environments. Among bacterial communities (Fig. 2B), cations (Ca and Mg) clustered with CEC and pH, while nutrient variables such as P, K, Na, NO_3_ ^-^, and OM formed a separate grouping. These groupings point in similar directions and indicate they have an influence on beta diversity in bacterial communities. For fungal communities (Fig. 2D), vectors were more uniformly oriented, indicating that the soil properties influenced fungal beta diversity in a more consistent manner.

**Figure 2.**
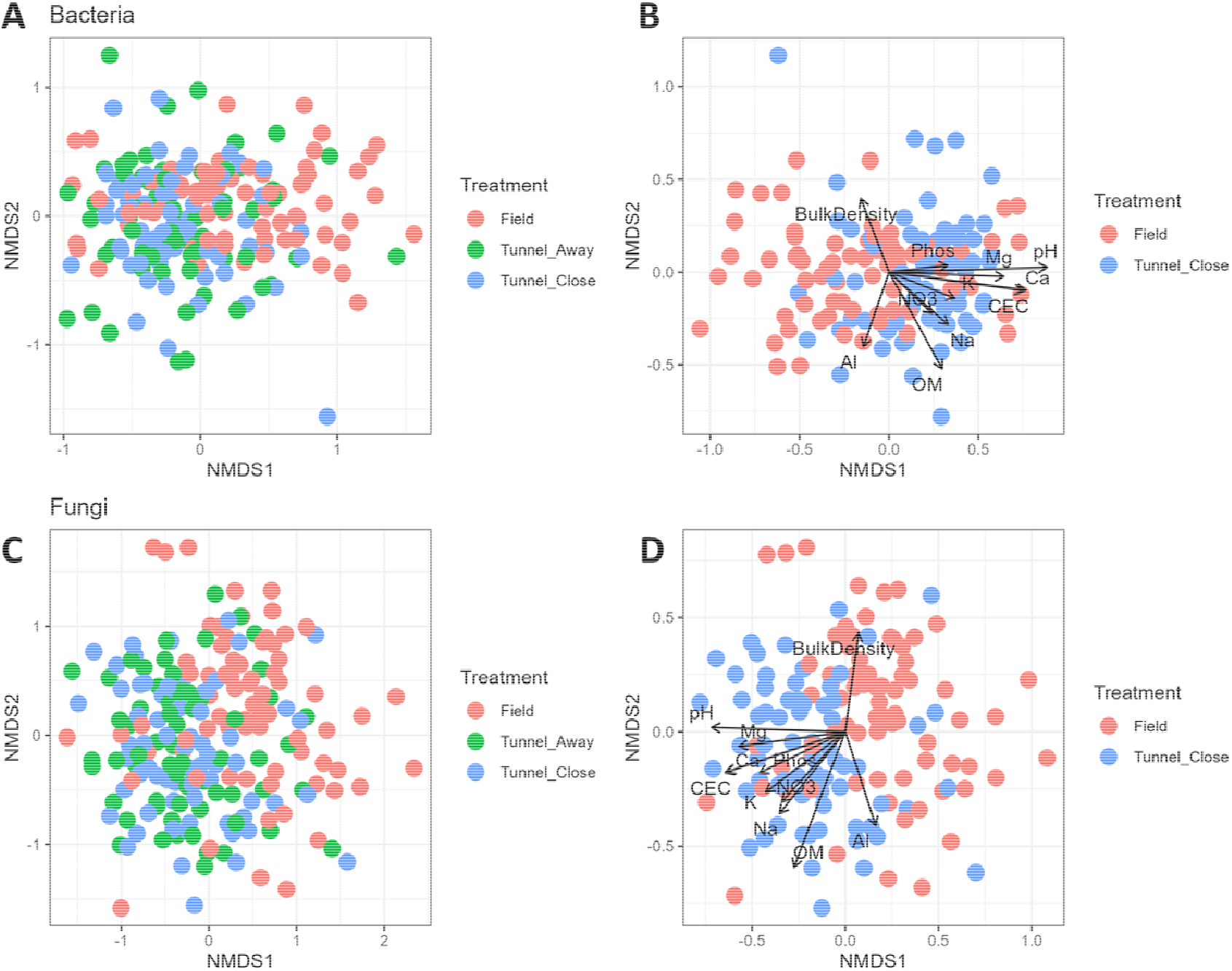
NMDS ordinations showing bacterial and fungal community □-diversity across field and high-tunnel environments. (A) Nonmetric multidimensional scaling (NMDS) ordination of bacterial communities among all environments. Samples are colored by environment and drip-line position: Field (red), Tunnel_Away (green), and Tunnel_Close (blue). (B) NMDS ordination of bacterial communities from Field (red) and Tunnel_Close (blue) samples with significant vectors of soil properties. (C) NMDS ordination of fungal communities, with samples colored by environment and dripline position: Field (red), Tunnel_Away (green), and Tunnel_Close (blue). (D) NMDS ordination of bacterial communities from Field (red) and Tunnel_Close (blue) samples with soil property vectors. Each arrow is labeled with its corresponding soil property abbreviation: bulk density, cation exchange capacity (CEC), pH, Mg, Ca, Al, NO_3_ ^-^, Na, K, and P.

### Composition of soil microbial communities in field and high-tunnel environments

Soil bacterial community composition differed between field and high-tunnel environments. Mean relative abundance profiles (Fig. 3A) showed that all environments were dominated by Halanaerobiaeota, Nitrospirota, and Actinobacteriota phyla. High-tunnel environments were significantly enriched in Firmicutes, Deinococcota, Patescibacteria, Halanaerobiaeota, and Halobacterota. Field environments were enriched in Acidobacteriota, Verrucomicrobiota, Myxococcota, and other phyla. There were no significant differences in bacterial phyla between locations close to and away from the driplines within high-tunnels (Table S1). Differential abundance analysis using ALDEx2 identified multiple phyla that showed significant differences between high-tunnel and field soils (Fig. 3B). Firmicutes, Deinococcota, and Halanaerobiaeota exhibited a relative increase in their abundance in high-tunnel soils, whereas Acidobacteriota, Verrucomicrobiota, Planctomycetota, Mycococcota, and several others decreased. At the genus level (Fig. 3C), many taxa showed strong responses: genera such as Longispora, Truepera, Paenisporosarcina, and Salinimicrobium increased in high-tunnel soils, whereas many others, including Bradyrhizobium, Haliangium, Massilia, Rhizobacter, and many others, decreased. These patterns suggest that nutrient levels under different management systems drive shifts in soil microbial communities and that certain bacterial groups are better adapted to drier or saline microclimates often associated with high-tunnel cultivation. A greater relative abundance of several phyla in field soils indicates a preference for lower nutrient levels or for more complex resource availability.

**Figure 3.**
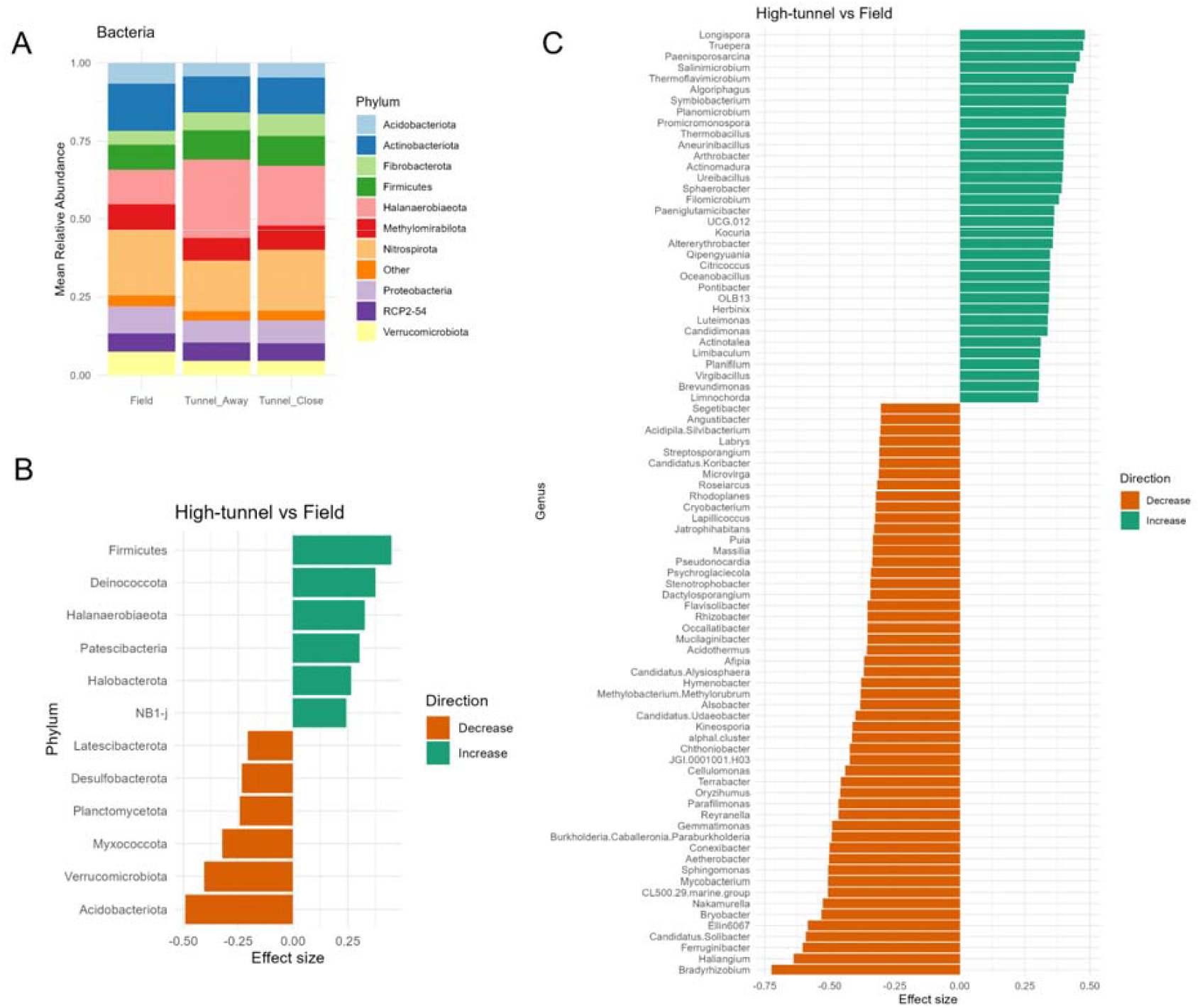
Relative abundance and differential enrichment of bacterial taxa in high tunnel vs. field soil samples. (A) Stacked bar plot showing mean relative abundance of bacterial phyla in field soils and high tunnel soils either away (Tunnel_Away) or close (Tunnel_Close) to the dripline. Bars are colored by phylum as indicated in the key on the right. (B) Differential abundance analysis of bacterial phyla using ALDEx2. Bars indicate the effect size, with green representing a relative increase and red representing a relative decrease in high tunnel soils compared to field soils. (C) Differential abundance analysis of bacterial genera using ALDEx2. Bars indicate the effect size, with green representing a relative increase and red representing a relative decrease in high tunnel soils compared to field soils.

Similarly, fungal soil community composition also differed between environmental types.

Mean relative abundance profiles (Fig. 4A) revealed that Mortierellomycota, followed by Ascomycota, Basidiomycota, and Sanchytriomycota, dominated fungal communities across all the environments. Differential abundance analysis using ALDEx2 showed that Ascomycota, Basidiomycota, Mortierellomycota, and Mucoromycota relatively increased in high tunnel soils, but Glomeromycota, Kickxellomycota, and Sanchytrioyctoa showed relative decreases (Fig. 4B).

**Figure 4.**
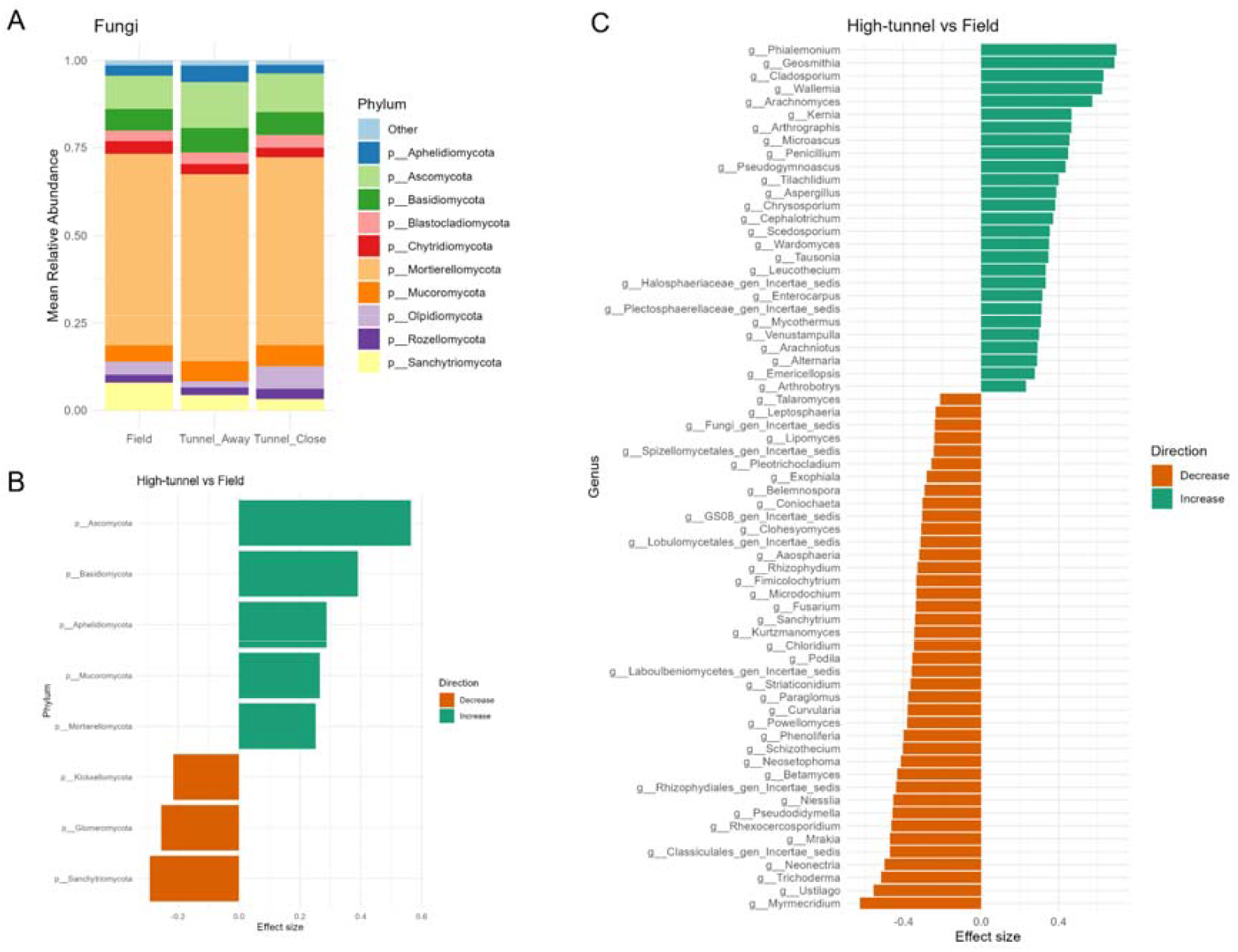
Relative abundance and differential enrichment of fungal taxa in high tunnel vs. field soil samples. (A) Stacked bar plot showing mean relative abundance of fungal phyla in field soils and high tunnel soils either away (Tunnel_Away) or close (Tunnel_Close) to the dripline. Bars are colored by phylum as indicated in the key on the right. (B) Differential abundance analysis of fungal phyla using ALDEx2. Bars indicate the effect size, with green representing a relative increase and red representing a relative decrease in high tunnel soils compared to field soils. (C) Differential abundance analysis of fungal genera using ALDEx2. Bars indicate the effect size, with green representing a relative increase and red representing a relative decrease in high tunnel soils compared to field soils. Only genera with effect sizes >|0.3| are presented.

At the genus level (Fig. 4C), genera such as *Phialemonium, Geosmithia, Cladosporium, Wallemia, Arachnomyces*, and *Kernia* showed relative increases in high tunnel soils. In contrast, a wide range of genera, including *Myrmecridium, Ustilago, Trichoderma, Neonectria*, and many others, relatively decreased. These shifts show that high-tunnel environments selectively favor fungal taxa adapted to altered microclimatic conditions and soil physicochemical parameters, resulting in increased relative abundances of stress-tolerant or disturbance-associated genera, while simultaneously reducing the prevalence of genera reliant on more heterogeneous or natural field soil conditions. Such selective enrichment or suppression at both the phylum and genus levels reflects the profound impact of cultivation practices and environmental modification on soil fungal community structure and potential ecosystem functioning.

No significant differences in relative abundances of fungal phyla or genera between within or away from driplines within high-tunnels (Table S1). Notably, the relative abundance of Glomeromycota, the phylum to which arbuscular mycorrhizal fungi (AMF) belong, was significantly negatively related to soil P content (Fig. 5; Spearman rho = -0.49, p<0.0001), suggesting that high P levels under HT conditions suppress the prevalence of these key plant symbionts. However, the correlation was not significant for AMF spore counts (Spearman rho = 0.14, p = 0.102), suggesting that AMF are present as spores, but do not form active symbiotic interactions in HTs under high P conditions.

**Figure 5.**
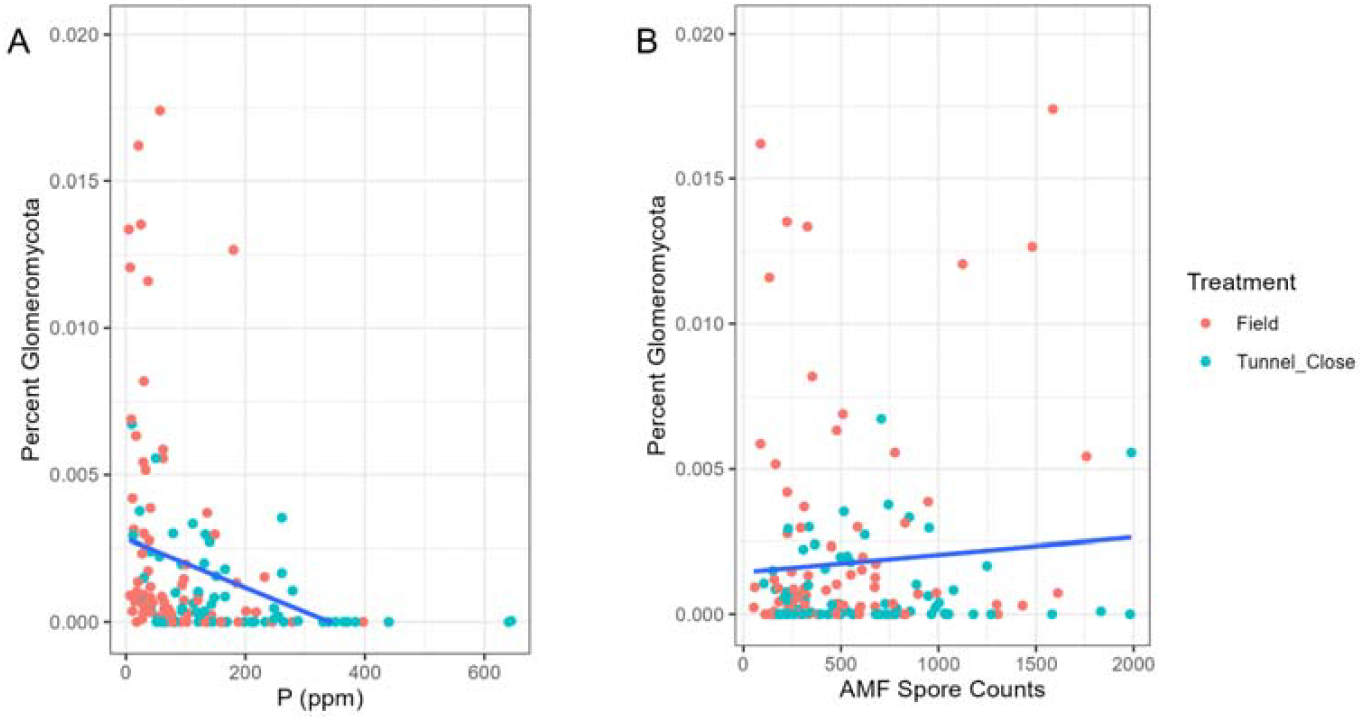
Correlation of percent Glomeromycota abundance with soil phosphorus and arbuscular mycorrhizal fungi (AMF) spore counts between environments. (A) Relationship between percent of Glomeromycota sequences and soil phosphorus (ppm). (B) Relationship between percent Glomeromycota sequences and AMF spore counts. Samples are colored by treatment, field (red), tunnel_close (blue), with a fitted regression line indicating the trend.

Glomeromycota abundance showed weak to moderate associations with soil phosphorus concentrations and AMF spore counts (Fig. 5). Glomeromycota was significantly negatively related to increasing phosphorus concentration (Spearman G = -0.49, p < 0.0001). This negative relationship was more pronounced in high-tunnel samples than in field samples. In contrast, the relationship between Glomeromycota abundance and AMF spore count was weak and not significant (Fig. 5B, Spearman G = 0.14, p = 0.102), with substantial variability among samples.

### Microbial co-occurrence networks among field and high-tunnel environments

Bacterial co-occurrence networks were distinct between field and high-tunnel environments (Jaccard similarity of 0.061 and 0.088 for field vs. away and close, respectively). Field bacterial networks were highly connected, with many high-degree nodes forming a dense central cluster (Fig. 6A, left), whereas tunnel-close and tunnel-away networks were less connected and comprised smaller, more dispersed clusters of co-occurring taxa (Fig. 6A, middle and right). Globally, field bacterial networks had more edges, were denser, and had higher average node degree than high-tunnel networks, and networks within and away from driplines were more similar to each other (Jaccard similarity of 0.294) than to field networks (Table 3).

**Table 3:**
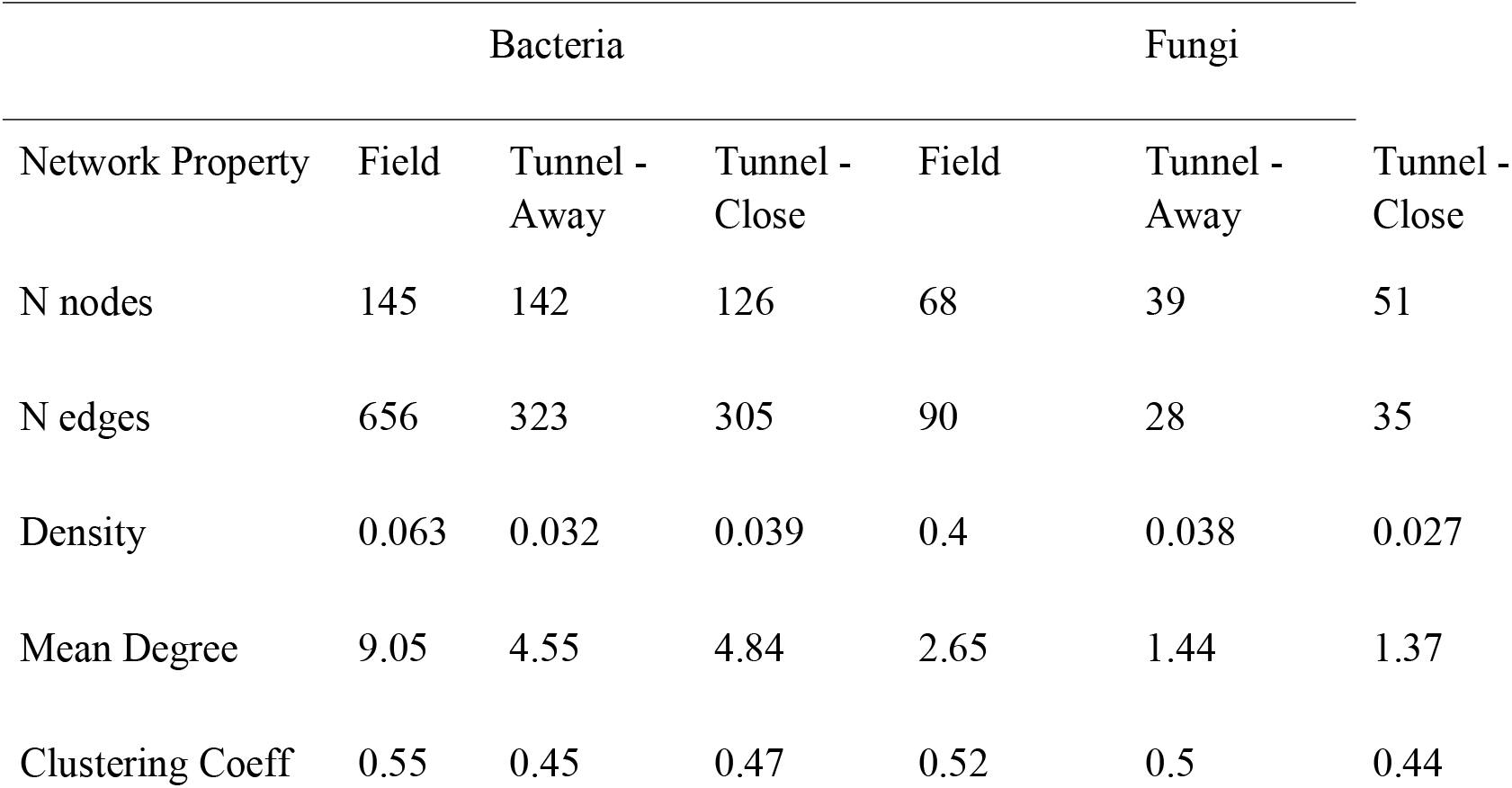
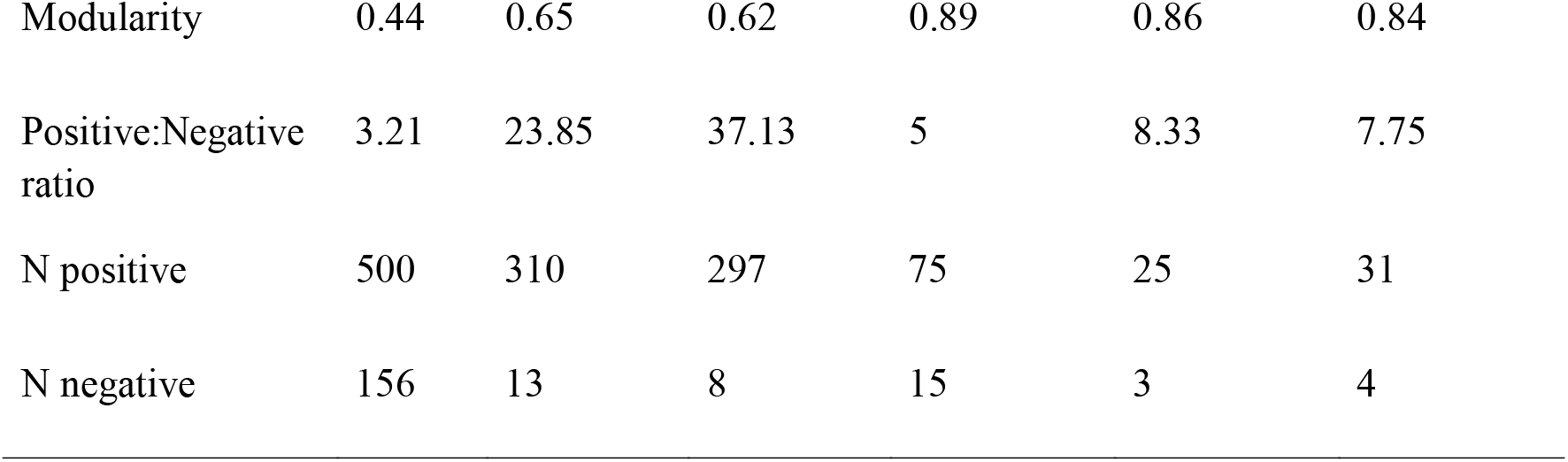
Co-occurrence network analysis.

**Figure 6.**
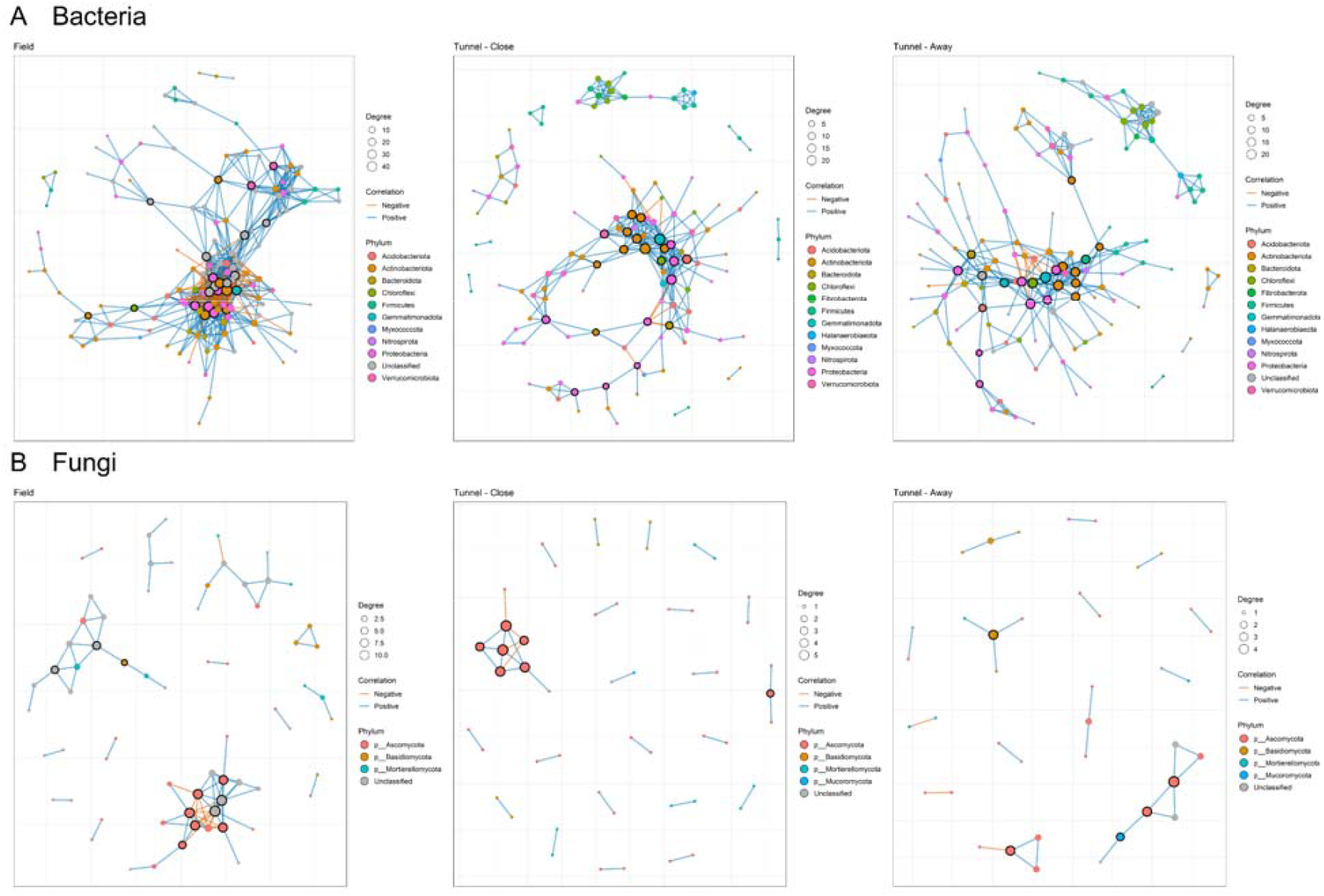
Microbial co-occurrence networks across field and high-tunnel environments. Bacterial (A) and fungal (B) co-occurrence networks are shown for field (left), tunnel - close to the dripline (middle), and tunnel - away from the dripline (right) environments. Nodes represent ASVs, colored by phylum and scaled by degree, while edges indicate positive (blue) or negative (red) significant correlations among taxa. Tightly-clustered nodes suggest greater interconnectivity among taxa.

Both high-tunnel bacterial networks also contained distinct modules of co-occurring taxa that were separated from the main network. Further, the ratio of positive: negative edges in the bacterial networks from high-tunnels was strikingly greater than that from fields. Fungal co-occurrence networks also differed between field and high-tunnel environments (Jaccard similarity of 0.068 and 0.088 for field vs. away and close, respectively), while being more similar within high tunnels (Jaccard similarity of 0.17 for away vs. close). Overall connectivity in fungal networks was lower than in bacterial networks, particularly under high-tunnel conditions. The field fungal network contained several distinct clusters of both positively and negatively associated taxa, whereas tunnel-close and tunnel-away fungal networks consisted mainly of small, low-degree modules with few edges (Fig. 6B). Across environments, field fungal networks had more nodes, more edges, and higher density than those from within high tunnels, whereas the ratios of positive to negative edges differed less among environments than observed for bacterial networks (Table 3).

## Discussion

This study provides a comprehensive comparison of soil microbial communities in high-tunnel and field environments across different locations in Minnesota, revealing significant differences in bacterial and fungal diversity, composition, and network structure. The results highlight the profound influence of management practices and soil physicochemical properties on microbial communities, with implications for sustainable agriculture and crop productivity.

### Microbial Diversity and Community Composition

High-tunnel soils exhibited higher levels of organic matter, pH, nitrate, phosphorus, and potassium than field soils, consistent with previous findings of nutrient accumulation in protected environments (Pierre et al. 2024). Despite these differences, alpha diversity metrics for both bacteria and fungi did not differ significantly between field and high-tunnel environments, suggesting that overall microbial richness and evenness are maintained across systems. However, beta diversity analyses revealed distinct community structures, with bacterial and fungal communities clustering separately between field and high-tunnel systems. This indicates that while overall diversity is similar, the specific taxa present and their relative abundances differ substantially between high-tunnels and fields. The increased abundance of the bacterial phylum Firmicutes in the soil may be influenced by the accumulation of salts and other nutrients, due to reduced nutrient leaching from relatively dry soil in the high-tunnels compared to field soil.

Previous studies show that high soil salinity increases the abundance of Firmicutes, which are Gram-positive and form spores, helping them resist salt stress (Schimel et al. 2007). Additionally, halophilic organisms, including Halanaerobiaeota and Halobacterota, were also enriched in the hypersaline soils of high-tunnels. Using the “salt-in” strategy, these bacteria break down complex organic matter via fermentative pathways, converting carbohydrates and other organic compounds into simpler fermentation products such as acetate, hydrogen, and organic acids (Roush et al. 2014). By producing these intermediary metabolites, Halanaerobiales play an ecological role in supporting the growth of sulfate-reducing and methanogenic microorganisms in saline conditions (Scheffer et al. 2025). Although high-tunnel soils are typically aerobic, temporary anaerobic conditions may arise due to poor drainage. Under such conditions, sulfate-reducing bacteria can acidify soil by forming sulfuric acid, altering nutrient availability, and affecting plant growth. Poor drainage combined with excessive organic matter can also lead to saturated soil for extended periods, creating anoxic microniches necessary for methanogenesis (Frey et al. 2011). The increased abundance of Deinococcota in high-tunnel soils likely reflects environmental stressors such as occasional extreme temperatures or dry periods during intermittent irrigation; however, their specific functional role is not well established (Kato et al. 2022). Similarly, Patescibacteria, generally found in wastewater treatment plants, would likely exist as part of complex microbial communities that facilitate nutrient flow via cooperative interactions in high-tunnel soils (Hu et al. 2024). Moreover, community composition analysis revealed increased abundance of heat- and salt-tolerant bacterial genera, including *Paenisporosarcina, Truepera, Thermoflavimicrobium, Thermobacillus, Salinimicrobium, Algoriphagus*, and others, in high-tunnel soils. While their ecological roles in high-tunnel soils remain hypothetical, their enrichment in these soils suggests that high levels of salt and organic matter, combined with dry, hot conditions, could impact soil functioning, decomposition, and nutrient cycling (Liang et al. 2022; Peng et al. 2024; Touzel et al. 2000).

Similarly, microclimates and soil fertility in high-tunnels altered soil fungal communities compared with open-field soils. The observed enrichment of Mortierellomycota, Ascomycota, and Basidiomycota across environments is consistent with global surveys showing that a small set of ascomycete and basidiomycete saprotrophs dominate many soils and respond strongly to shifts in moisture and nutrient status. The higher relative abundance of Ascomycota, Basidiomycota, Mortierellomycota, and Mucoromycota in high-tunnel soils suggests that these structures favor fast-growing, disturbance-or fertilizer-adapted taxa able to exploit elevated nutrient availability and more stable, but often drier or more thermally variable, microclimates (Baldrian et al. 2022; Yang et al. 2022).

The relative decline of Glomeromycota, Kickxellomycota, and Sanchytriomycota indicates that high tunnel conditions disfavor some more specialized or microhabitat-sensitive groups, particularly obligate symbionts such as AMF. This pattern aligns with broader evidence that intensive management, including fertilizer inputs and altered water regimes, often reduces the abundance and diversity of mycorrhizal fungi while favoring saprotrophic guilds that can capitalize on increased organic inputs and labile nutrients (Luo et al. 2021; Oliveira et al. 2024).

The observed increases in *Phialemonium, Geosmithia, Cladosporium, Wallemia, Arachnomyces*, and *Kernia* genera in high tunnel soil samples collectively signal a robust shift toward saprophytic fungal dominance, potentially driven by shared intensive management factors like frequent organic amendments (high levels observed in the high tunnels), moisture/humidity, and pH rises (often 7-9) that favor decomposers over competitors. These fungi, primarily keratinophilic (*Arachnomyces, Kernia*) or xerophilic/halotolerant (*Wallemia*) saprotrophs, alongside generalists (*Phialemonium, Geosmithia, Cladosporium*), thrive on decaying plant residues, composts, or keratin inputs, enhancing organic matter breakdown, nutrient cycling, and microbial diversity for improved soil fertility and plant access to resources (Rivera-Vega et al. 2022; Das et al. 2019; Williams and Ginzel 2022; Batson et al. 2022). While most indicate neutral-to-beneficial dynamics aligned with sustainable high-tunnel practices (e.g., *Phialemonium* and *Arachnomyces*), *Cladosporium* and *Geosmithia* diverge slightly due to potential disease risks, such as foliar leaf spots from humidity or insect-vectored diseases (Batson et al. 2022; Williams and Ginzel 2022). The mycotoxin potential of *Wallemia* during biomass buildup can be concerning. The relative decrease in diverse fungal genera like *Myrmecridium, Ustilago, Trichoderma, Neonectria*, and others in high tunnel soil samples, contrasted with a rise in saprophytic dominants (*Phialemonium, Geosmithia*), reflects a microbiome restructuring under intensive cropping, where organic amendments may have suppressed competitive or specialized fungi in favor of resilient decomposers. *Trichoderma*, a key biocontrol agent against root pathogens, often declines under suboptimal conditions, such as very dry or cold soil or non-ideal pH levels, giving better-adapted saprotrophs a competitive advantage (Kredics et al. 2003). Moreover, targeted irrigation in high-tunnels keeps the plant foliage and surface soil drier compared to open fields exposed to natural rainfall. Since wind-borne spores need moisture on the plant surface to germinate and infect, a lack of moisture may have reduced the abundance of *Ustilago* and *Neonectria* genera. The physical structure of a high-tunnel acts as a barrier, limiting air currents and thus reducing the natural wind dispersal of *Ustilago* teliospores from infected to healthy plants (Pataky and Snetselaar, 2006; Beresford and Kim, 2011). *Myrmecridium*, typically keratinolytic or oligotrophic, resembles *Kernia*/*Arachnomyces* patterns but declines under dry and hot conditions, favoring broader saprotrophs, indicating niche displacement rather than stress (Sun et al. 2022).

Soil nutrient levels, particularly phosphorus, nitrate, and potassium, were strongly correlated with microbial community composition. Notably, the relative abundance of Glomeromycota, the phylum containing arbuscular mycorrhizal fungi (AMF), was negatively correlated with soil phosphorus, suggesting that high nutrient conditions in high tunnels may suppress the prevalence of these beneficial symbionts. This finding aligns with the well-documented phenomenon that elevated phosphorus can inhibit AMF colonization and activity, potentially reducing the benefits of mycorrhizal associations for plant health and nutrient uptake (Balzergue et al. 2013).

Interestingly, AMF spore counts did not show the same negative correlation, indicating that while spores may persist, active colonization may be limited under high nutrient conditions. No significant differences in communities between high-tunnels were observed, suggesting that communities are relatively similar within HT environments. However, more information on soil properties, sampling timing, etc, is needed.

### Microbial Network Structure

Co-occurrence network analyses revealed that bacterial and fungal communities in high tunnels form distinct modules and exhibit different interaction patterns compared to field environments. Bacterial networks in high-tunnels were less dense and more modular, with particular clusters of taxa such as Chloroflexi and Firmicutes. These shifts in network structure suggest that microbial interactions and potential functional redundancy may be altered in high tunnels, potentially affecting ecosystem resilience and nutrient cycling. In contrast, fungal networks showed greater resilience, with less pronounced differences in network properties between field and high tunnel environments. Notably, microbial co-occurrence networks, especially those among bacterial taxa, had strikingly greater positive:negative edge ratios in high-tunnel versus field environments. This finding aligns well with the stress-gradient hypothesis, which predicts that facilitative species interactions are more common than competitive interactions under stressful environmental conditions (Harris and Bennett 2025). Key microbial stressors, such as drought and salinity, have been previously found to disrupt microbial networks, select for relatively more positive co-associations, and have a greater impact on bacterial versus fungal communities (Gao et al. 2022; Li et al. 2024). Together, this suggests that environmental stressors in high-tunnel environments generate more cooperative microbial communities, which may impact critical functions such as resource acquisition and detoxification of toxins or pollutants (Harris and Bennet, 2025).

Production in high-tunnels homogenizes soil microbial communities by imposing uniform microclimates and management practices, organic amendments, and targeted irrigation, which minimize environmental variability across Minnesota sites, leading to consistent nutrient accumulation (higher organic matter, pH, nitrate, phosphorus, potassium) and favoring salt/drought-tolerant bacteria and fast-growing saprotrophs. This results in tight beta-diversity clustering within high-tunnels, contrasting with heterogeneous field communities shaped by variable rainfall and leaching, while suppressing specialized genera. Operational timing could further drive differentiation: early-season nutrient spikes may select for copiotrophs, mid-season dryness may enrich stress-tolerant taxa, and asynchronous cycles or sampling relative to pre-vs. post-harvest could explain AMF spore abundance. Overall, the findings underscore the importance of considering management practices and soil properties when adopting sustainable agricultural systems. High-tunnels, while beneficial for extending growing seasons and protecting crops, may alter soil microbial communities in ways that could impact plant health and soil fertility. Specifically, the suppression of AMF under high-nutrient conditions suggests the need for careful nutrient management to maintain beneficial microbial associations.

Additionally, the distinct microbial network structures in high-tunnels highlight the potential for novel microbial interactions and functions, warranting further investigation. Future research should focus on elucidating the functional implications of these microbial community shifts, particularly in relation to crop productivity and disease resistance. Long-term monitoring of microbial communities under different management practices and nutrient regimes will be essential for developing strategies to optimize soil health and sustainability in both high-tunnel and field systems. Furthermore, exploring the potential of microbial inoculants or tailored management practices to enhance beneficial microbial associations in high-tunnels could open new avenues to improve crop yields and resilience.

## Acknowledgements

We want to thank all the extension educators of Minnesota who collected soil samples for AMF spore quantification and soil microbiome analyses. Mention of trade names or commercial products in this publication is solely for the purpose of providing specific information and does not imply recommendation or endorsement by the U.S. Department of Agriculture (USDA). USDA is an equal opportunity provider and employer.

## Author Contributions

**MT -** Writing – review and editing, Methodology, Writing – original draft, Conceptualization, Investigation, Formal analysis, Validation. **DS -** Formal analysis, Writing - review and editing. **NH -** review and editing. **DK -** Conceptualization, Methodology, Investigation, Writing – review and editing

---

**Supplementary Table 1:**
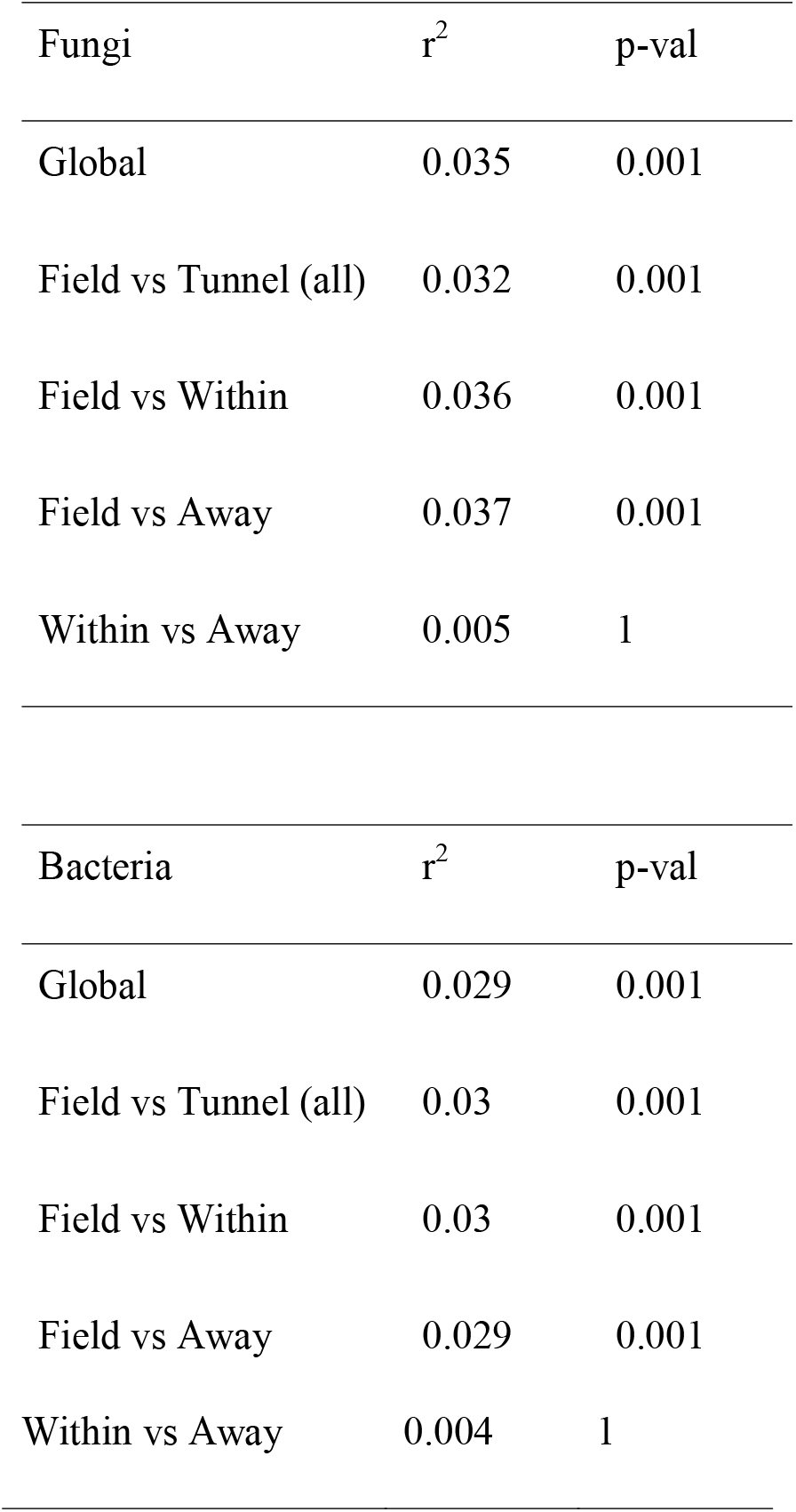
PERMANOVA analysis of fungal and bacterial communities among environments.

